# Modern Machine Learning: Partition & Vote

**DOI:** 10.1101/2020.04.29.068460

**Authors:** Carey E. Priebe, Joshua T. Vogelstein, Florian Engert, Christopher M. White

**Affiliations:** Johns Hopkins University; Harvard University; Microsoft Research

**Keywords:** Decision Forests, Deep Networks, Classification, Brain functioning

## Abstract

We present modern machine learning, focusing on the state-of-the-art classification methods of decision forests and deep networks, as partition and vote schemes. This illustrative presentation allows for both a unified basic understanding of how these methods work from the perspective of classical statistical pattern recognition as well as useful basic insight into their relationship with each other … and potentially with brain functioning.

## 1 Introduction

The classical statistical formulation of the classification problem consists of

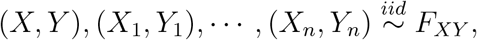

where 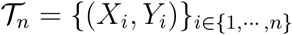 is the training data and (*X, Y*) represents the to-be-classified test observation *X* with true-but-unobserved class label *Y*. We consider the simplest setting in which *X* is a feature vector in 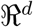 and *Y* is a class label in {0, 1}. The goal is to learn a classification rule 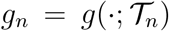 mapping feature vectors to class labels such that the probability of misclassification 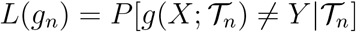 is small.

Stone’s Theorem for universally consistent classification (Stone, 1977; Devroye et al., 1997) demonstrates, loosely speaking, that a successful classifier can be constructed by first partitioning the input space into cells – the partition depends on *n* – such that the number of training data points in each cell goes to infinity but slowly in *n*, and then estimating the posterior *η*(*x*) = *P*[*Y* = 1|*X* = *x*] locally by voting based on the training class labels associated with the training feature vectors in cell 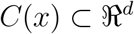 in which the test observation falls. That is, letting *S*(*x*) = {*X*_1_, … , *X*_*n*_} ∩ *C*(*x*) be the subset of *X*_*i*_’s falling in cell *C*(*x*), and letting *N*(*x*) = |*S*(*x*)| = Σ_*i*_ *I*{*X*_*i*_ ∈ *C*(*x*)} be the cardinality of this set, we consider the posterior estimate 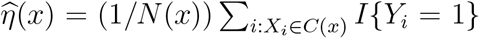. Then, under some technical conditions on the manner of choosing the sequence of partitions 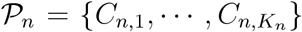, the plug-in rule 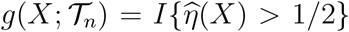 is universally consistent: *L*(*g*_*n*_) → *L*^*^ almost surely for any *F*_*XY*_, where *L*^*^ is the Bayes optimal probability of misclassification.

## 2 Decision Forests & Deep Networks

In the context of our formulation of the classification problem, we provide a unified description of the two dominant methods in modern machine learning, the decision forest (DF) and the deep network (DN), as ensemble partition and vote schemes. This allows for useful basic insight into their relationship with each other … and potentially with brain functioning.

### 2.1 Decision Forests

Decision forests, including random forests and gradient boosting trees, demonstrate state-of-the-art performance in a variety of machine learning settings. DFs have typically been implemented as ensembles of axis-aligned decision trees – trees that split along canonical feature dimensions only – but modern extensions employ axis-oblique splits (Breiman, 2001; Criminisi and Shotton, 2013; Athey et al., 2019; Tomita et al., 2020).

Given the training data 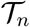, each tree *t* in a random forest constructs a partition 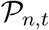 by successively splitting the input space based on a random subset of the data (per tree) and then choosing a hyperplane split based on a random subset of the dimensions (per node) using some criterion for split utility at each node such as maximal reduction in class confusion for the training data falling to that node. The details provide ample fodder for myriad implementations, and a simple depiction necessarily suffers from an inability to Visualize input space 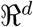 for *d* > 2, but Figure 1 illustrates the idea.

**Figure 1:**
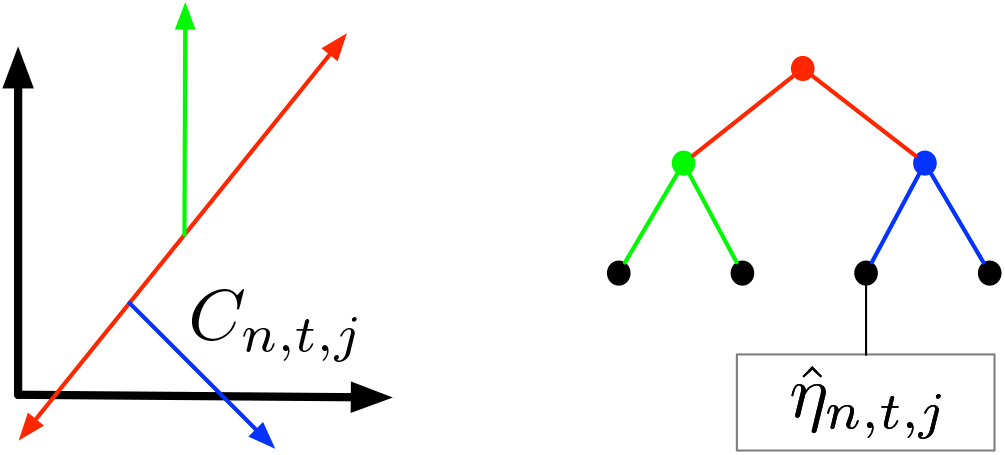
A tree in the forest. Given the random subset 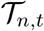 of the training data allocated to tree *t*, the root node (red) performs a hyperplane split of 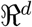 based on a random subset of dimensions; the two daughter nodes (green and blue) split their respective partition cells based on a separate random subset of dimensions allocated to each node; etc. In the end, this tree results in a partition of 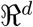 with the leaf nodes corresponding to partition cells for which training data class labels yield local posterior estimates. The forest classifies the test observation feature vector *X* by voting over trees using the cells *C*_*n,t*_(*X*) in which *X* falls.

The depth of each tree is a function of *n* and involves a tradeoff between leaf purity and regularization to alleviate overfitting. Details aside, each tree results in a partition 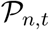 of 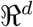 and each partition cell – each leaf of each tree – admits a posterior estimate 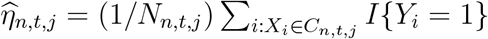 based on the class labels of the training data feature vectors that fall into cell *C*_*n,t,j*_.

The conceit of the forest is that of resampling: by choosing a separate subset 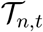 of the training data for each tree *t*, the ensemble forest posterior estimate is superior to that of any individual tree. Regardless, the overall classifier is seen to be a partition and vote scheme, and under appropriate conditions on the manner of choosing the partitions 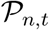 we have 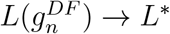

### 2.2 Deep Networks

Deep networks are extraordinarily popular and successful in modern machine learning. (LeCun et al., 2015; Sze et al., 2017; Montúfar et al., 2014; Montúfar, 2017). As depicted in Figure 2, each internal node *υ*_*ℓ,k*_ in layer *ℓ* of the network gathers inputs 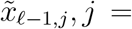 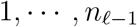 from the previous layer, weighted by *w*_*ℓ*−1*,j,k*_, *j* = 1, … , *n*_*ℓ*−1_, and outputs

**Figure 2:**
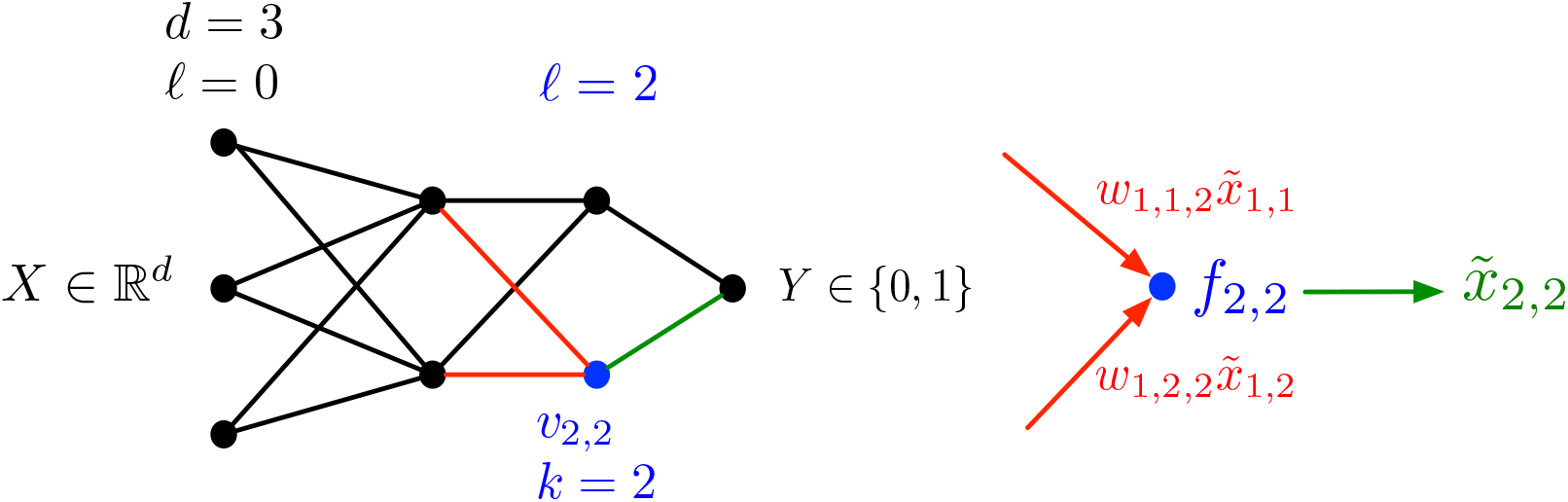
A (shallow) deep network. Given the training data 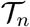, the 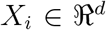 are passed through the network. At layer *ℓ* and node *υ*_*ℓ,k*_ (the blue node is *υ*_2,2_) the inputs 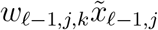 (red) are transformed via hyperplane activation function *f*_*ℓ,k*_ and output as 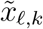 (green). Thus node *υ*_2,2_ receives non-zero input 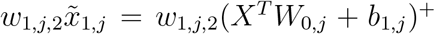 from node *υ*_1,*j*_ if and only if the linear combination of the multivariate *X*, *X*^*T*^*W*_0,*j*_, is on the preferred side of hyperplane defined by *f*_1,*j*_ (and weight *w*_1,*j*,2_ is non-zero). The output of node *υ*_2,2_ is 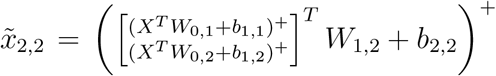; that is, *υ*_2,2_ provides a further hyperplane refinement of 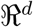.

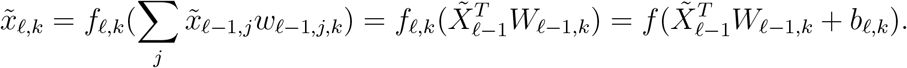

When using the so called ReLU (rectified linear unit) function max(0, ·) = (·)^+^ as the activation function *f*, each node performs a hyperplane split based on a linear combination of its inputs, passing a non-zero value forward if the input is on the preferred side of the hyperplane; data in the cell defined by the collection of polytopes induced by nodes {*υ*_*ℓ*−1,*j*_}, weighted, falls into node *υ*_*ℓ,k*_ and is output based on a partition refinement. Letting 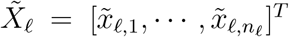 be the output of layer *ℓ* and 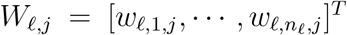 be the weights from layer *ℓ* to the *j*th node in layer *ℓ* + 1, node *υ*_*ℓ*+1,*j*_ sees 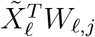 and outputs 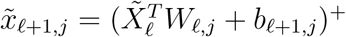. Thus a node in the last internal layer corresponds to a union of hyperplane-induced partition cells, defined via composition of all the nodes earlier in the network.

For conceptual unification with DFs, consider passing all the training data through the network; the collection of *X*_*i*_’s falling into each of the nodes in the penultimate layer, together with their class labels, induces local posterior estimates for each of these cells. As with a single tree in a DF wherein the test observation feature vector *X*, suitably transformed, falls into one and only one leaf partition cell, Montúfar et al. (2014) argues that DNs with ReLu activation functions fold space such that a subset of 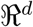 maps to exactly one fold. Thus for a DN the input *X* falls into a final network partition cell in the last internal layer. Unlike in DFs, for a DN the input *X* activating a node in the penultimate layer does not uniquely specify which partition cell *X* falls into; rather, it merely indicates that *X* falls into the set of partition cells corresponding to that node. For this reason, whereas in a DF each *X* activates only a single leaf node, in a DN each *X* can activate many (even all) cells in the penultimate layer, though with different activation energies. Thus while for a DF the ensemble is realized by voting over a forest of trees, for a DN we have membership in a single final network partition cell. On the other hand, each cell in a DF is constructed via a simple local splitting process, while for a DN a complex parameter estimation solution is employed to tailor the individual final network partition cells to the training data. In other words, both DFs and DNs can be seen to use the same representation space, though they achieve their particular representation via different estimation (“learning”) algorithms.

The parameters for this partition and vote scheme are estimated (“learned”) so as to make the network input/output fit the training data 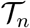, with much current effort involving aspects of regularization to alleviate overfitting and capacity as a function of the number of layers and nodes per layer.

## 3 An Illustrative Comparative Example

Practical application of both DFs and DNs involve a multitude of design and computational considerations; Figure 3 presents a simple illustration of comparative performance. The experimental setup is as follows (see Park (2020) for full reproduction capability). We choose two CIFAR-10 classes (Krizhevsky, 2012, 2015). There are 6000 images per class, which are split into training (5000 per class) and testing (1000 per class). For DF we use the R package “ranger” (Wright and Ziegler, 2017); for DN we use the R package “keras” (Chollet et al., 2015). The figure shows classification performance *L*(*g*) against the number of training samples *n*.

**Figure 3:**
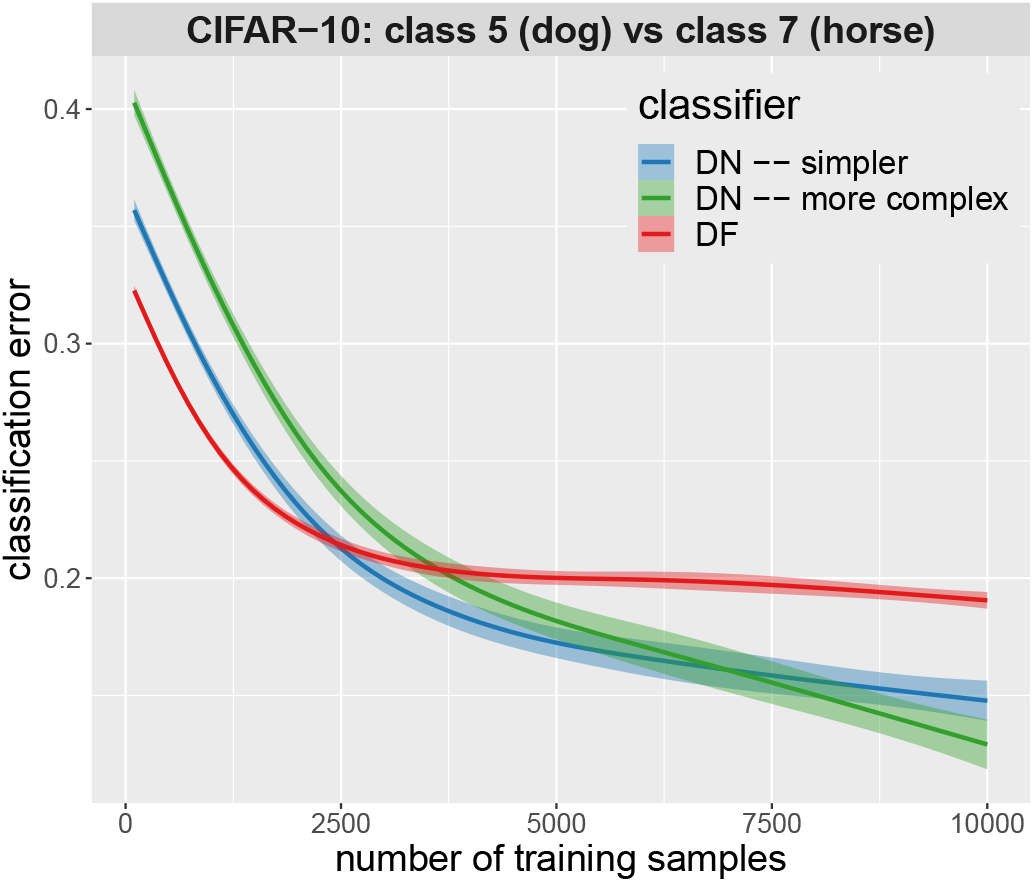
Performance comparison of a DF to two DNs: DF in red; DN in green and blue, with blue the simpler architecture and green the more complex. The x-axis is training set sample size *n*; the y-axis is classification error *L*(*g*).

DFs, with the number of trees and their depths going to infinity as *n* → ∞, are inherently nonparametric. DNs, with a fixed architecture, are parametric, albeit with a large enough number of parameters, and a mysterious enough implicit regularization effect, to blur this distinction in many settings. The simple comparative example presented in Figure 3 demonstrates the expected DN bias-variance tradeoff: a simpler DN outperforms a more complex DN when *n* is small, with this relationship switched for *n* large enough. DF (properly designed and relaxed in *n*) will eventually outperform any fixed-architecture DN as *n* increases; in this example the maximum *n* considered is too small for this limiting effect to obtain. For *n* very small, DF superiority is due to the fact that the variance of the DN that is required to get good performance for larger *n* renders its risk worse at low *n*.

## 4 A relationship with brain functioning

Learning in both DNs and DFs can be thought of as ensemble partition and vote functions implemented by a network of nodes. Intriguingly, learning in biological brains can be viewed similarly. In brains, a ‘node’ can correspond to a unit across many scales, ranging from synaptic channels (which can be selectively activated or deactivated due to the synapses’ local history), to cellular compartments, individual cells, or cellular ensembles (Vogelstein et al., 2019). At each scale, brains learn by partitioning feature space. The feature space that is partitioned is the set of all possible sensory inputs; a ‘part’ corresponds to a subset of ‘nodes’ that tend to respond to any given input. An example is the selective response properties that define cortical columns–columns in the sensory cortex (Mountcastle, 1997).

Brains also vote, where voting is a pattern of responses based on neural activation that indicate which stimulus evoked a response (Machens et al., 2005).

Figure 4 illustrates neural selectivity to features of sensory input in a larval zebrafish, and indicates a learned partition and vote framework for brain functioning (Naumann et al., 2016). Specifically, this image shows all motion-sensitive nodes (*n* = 76, 604) in a brain. Each node is a dot, color coded for preferred motion direction. The image indicates that each node’s activity indeed corresponds to a partition of feature space, which is actively refined through sensory experience and learning. A relationship with DFs and DNs, at least at a basic level, is clear. The utility of this relationship, for either machine learning or neuroscience, is the subject of endless conjecture and refutation.

**Figure 4:**
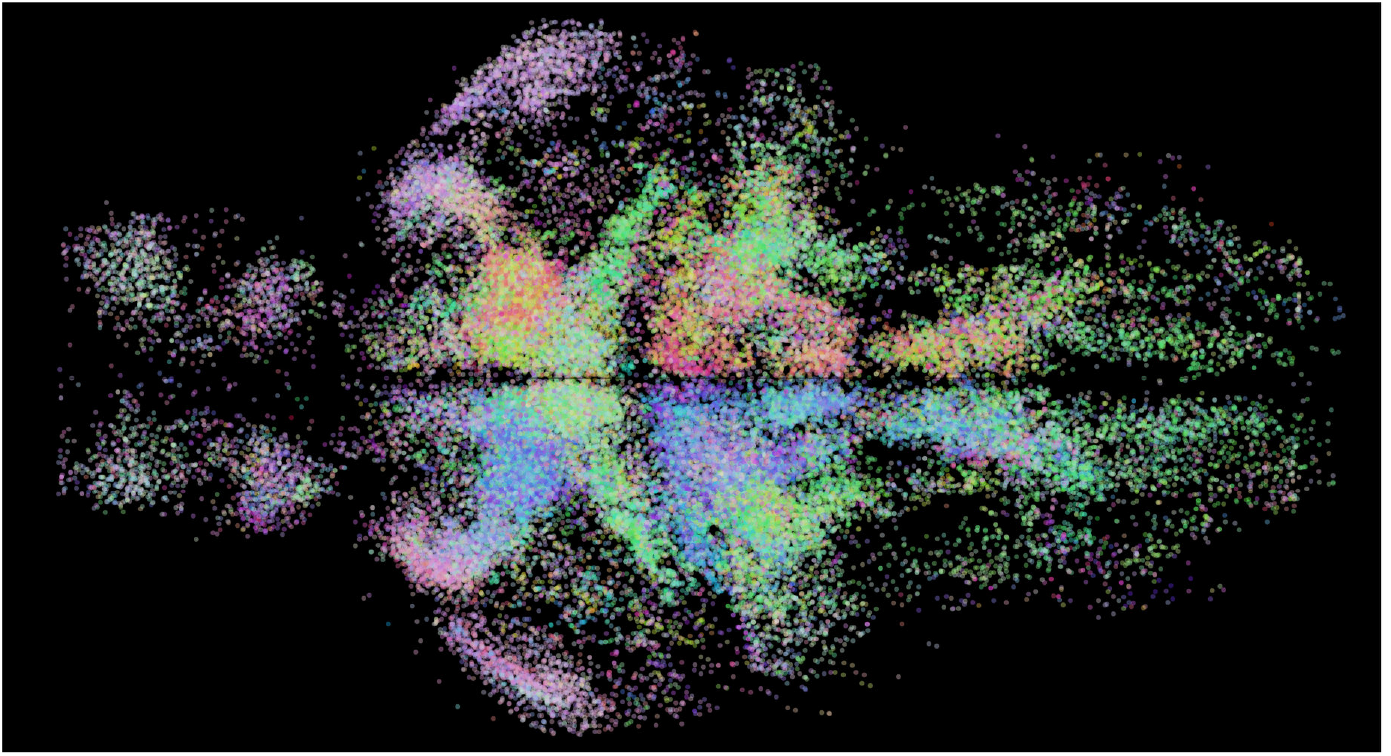
Whole-brain activity maps reveal processing stages underlying the optomotor response.

## 5 Conclusion

“Many important classification rules partition 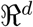 into disjoint cells *A*_1_, *A*_2_, … and classify in each cell according to the majority vote among the labels of the *X*_*i*_’s falling in the same cell” (Devroye et al. (1997), p. 94). The purpose of this short note is to describe state-of-the-art machine learning methods simplified in classical statistical pattern recognition terms. The depiction of deep networks and decision forests as partition and vote schemes allows for both a unified basic understanding of how these methods work from the perspective of classical classification as well as useful basic insight into their relationship with each other … and potentially with brain functioning. Elaborate generalization of the partition and vote framework is the subject of intense research – theoretical, methodological, and for myriad applications – but the fundamental idea is here presented in its most basic form.

The representational equivalence presented herein may enable ‘porting’ theoretical and conceptual advances between decision tree/forest approaches and deep network approaches. For example, nonparametric neural networks were recently introduced (Philipp and Carbonell, 2017) as a way to dynamically grow networks with larger sample sizes. While interesting, this approach lacks any formal statistical guarantees, such as those enjoyed by DFs. While the theory for DNs include universal consistency under certain conditions for network size growing, there are no algorithms (to our knowledge) that actually implement such an approach in a certifiably consistent fashion. Perhaps universal consistency proofs are available for these kinds of networks, if one more carefully investigates the relationships between DFs and DNs. Similarly, the double descent phenomenon has recently gained much attention in the DN world (Belkin et al., 2019). This effect is well-known and understood in the DF world: a single tree will interpolate and therefore overfit, yielding zero training error. But when simply adding more random trees, training error of course cannot decrease, but generalization error will continue to decrease. Finally, an open question about DFs is how to achieve parametric convergence rates. Empirically, DNs seem to converge faster. Perhaps this is due to the existence of a certain kind of ‘lateral connections’, which are not present in trees, but are present in directed acyclic networks (such as feedfoward deep networks). Perhaps the edges linking nodes in a DF that generalize it from being a pure tree provide the additional structure needed to regularize more effectively for finite samples, thereby improving convergence rates.

What does all of this mean with respect to biological, rather than artificial, learning? A large community of AI experts currently believe that DNs are a good model of brains. This consists of two ingredients: the learning algorithm and the representation space. While it remains hotly contested whether back propagation is a reasonable model of learning in the brain, as illustrated above there is little doubt in the community regarding whether brains partition feature space. In that sense, perhaps modeling the brain as a partition and vote machine, rather than as a deep learning machine, could alleviate some of the limitations of the current model. One specific claim this model would make, which does not require any statement on the details of the learning algorithm, is that the brain operates more like a kernel learning machine (which makes decisions using only ‘local’ information in feature space) than a polynomial regression machine (which makes decisions more ‘globally’). Another nice property of the model of the brain as a partition and vote learning machine is that it naturally makes predictions about both behavior and brain activity. Specifically, if one could design an experiment to identify the boundaries of the partitions for a given ensemble of neurons, one would predict that perturbing the stimulus such that a different ensemble of neurons is activated would lead to behavioral changes, whereas perturbing the stimulus by the same amount such that it stays within the partition would not lead to behavioral changes. This is in contrast to other large scale models of the brain, such as the free energy principle, which under certain assumptions makes predictions about behavior, but without a clear tie to the neural implementation.

